# AIPID: MAD-ML-Powered AIP Discovery Platform

**DOI:** 10.1101/2025.07.14.664584

**Authors:** Ananya Anurag Anand, Kaushik Urkude, Sintu Kumar Samanta

**Affiliations:** Department of Applied Sciences, Indian Institute of Information Technology Allahabad, Prayagraj 211012, Uttar Pradesh, India

**Keywords:** Peptides, Anti-inflammatory, Inflammation, Machine Learning, Random Forest, AIPID

## Abstract

Inflammation is a biological defense mechanism against harmful stimuli such as infection, tissue injury, or toxic agents, which, if prolonged, can lead to chronic inflammatory disorders. The limited safety and tolerability of current anti-inflammatory drugs emphasize the need for novel, selective therapeutic agents. Anti-inflammatory peptides (AIPs) have emerged as promising candidates owing to their ability to selectively target diseased cells while sparing healthy tissue. However, the identification of AIPs remains constrained by labor-intensive and expensive wet-laboratory screenings. To address this challenge, we have developed AIPID, an interactive, publicly accessible web-application for faster identification of anti-inflammatory peptides. Central to our platform is our Motif-Analysis-Driven Machine Learning (MAD-ML) model, based on the approach of representative negative dataset selection, combining iterative random sampling with motif profiling to enhance dataset diversity and model robustness. Finally, AIPID employs a Random Forest-based classifier trained on motif-filtered, biologically relevant peptide sequences, classifying inputs as AIPs or non-AIPs based on sequence-derived physiochemical descriptors. The model demonstrated excellent performance, achieving sensitivity of 95.16%, specificity of 99.98%, and F1 score of 98.14%, outperforming existing models and correctly predicting 18 of 19 experimentally validated AIPs. The AIPID platform offers an intuitive, multi-page interface for sequence-based peptide prediction, exploration of UniProt-derived AIP repositories, and access to statistical insights on peptide properties. The application is freely available at https://aipid-app-version1.streamlit.app/, providing a valuable resource for the peptide therapeutics research community.

## 1. Introduction

Inflammation serves as a defense mechanism for higher organisms against pathogenic agents and infections **(Khan et al., 2023).** Inflammatory responses occur when tissues are harmed under normal conditions by trauma, toxins, disease, or heat. Inflammation can be acute or chronic **(Deng et al., 2022).**

Acute inflammation is a type of innate immunological response, while chronic inflammation lasts for a long time and leads to a number of debilitating chronic diseases, including cancer, autoimmune disorders, cardiovascular disease, and neurological diseases.

Statistics reveal that, nearly three of five people die due to chronic inflammatory all over the world (Deng et al., 2022; Barcelos et al., 2019; Deepak et al., 2019;). Undoubtedly, chronic inflammation has posed a significant risk to human well-being. Glucocorticoids, nonsteroidal anti-inflammatory medications (NSAIDs), and some biologicals are the mainstays of treatment for autoimmune illnesses and chronic inflammation (Bindu et al., 2020). However, these have multiple adverse effects due to low target-specificity (Dendoncker and Libert, 2017). Additionally, the body develops resistance against many of these in the long-term. Thus, to overcome the two major problems of low target-specificity and resistance-induction, there is an urgent requirement for the identification or rational design of novel anti-inflammatory agents.

Peptide therapeutics, such as anticancer peptides and antibacterial peptides, have attracted a lot of attention in recent years due to their attractive advantages including efficacy, safety, ease of synthesis, and high selectivity (Deng et al., 2022). Anti-inflammatory peptides (AIPs) are a type of therapeutic peptides that exhibits anti-inflammatory properties. Generally, AIPs are short linear peptides composed of 10–50 amino acids, sometimes upto 120 amino acids. AIPs work by preventing excessive pro-inflammatory reactions, promoting anti-inflammatory responses, and modifying immune cell differentiation (Sun et al., 2018; Heinbockel et al., 2021).

As of now, the majority of AIPs that have been discovered are either endogenous peptides or obtained from natural sources. Examples of these include the sea snake-derived hydrostatin-SN1, melittin from bee venom, and endogenous neuropeptide vasoactive intestinal peptide (Zhang et al., 2020; Jiang et al., 2016; Lee et al., 2014). Some synthetic peptides have also been shown to inhibit inflammatory responses, such as, the BCL-3-mimetic (Collins et al., 2015; Usmani et al., 2017). These studies indicate that AIPs have great potential to be a new alternative therapy for treating inflammation.

Discovering AIPs against several autoimmune disorders and inflammatory diseases involves rigorous wet lab procedures, in addition to time and funds to identify them. Therefore, faster prediction of potential AIPs necessitates the development of an effective ML model. A few computational models have been developed for predicting AIPs, but lack desired accuracy and specificity, the reason being that most of these utilize the same and a small number of sequences, which limits the performance of the model. Additionally, most of the existing AIP-prediction models only take into consideration a single approach, without innovating on them. Numerous studies have demonstrated that the ensemble learning model performs better than the model based on a single algorithm (Deng et al., 2022; Jiang et al., 2021; Mishra et al., 2019). Although it was believed that using an ensemble learning strategy would greatly enhance the performance of AIP identification, despite being more efficient they did not demonstrate a significant improvement. Two ensemble learning methods—PreTP-EL and PreTP-Stack—have recently been reported, but their potential for AIP prediction remains limited. A sequence-based stacking ensemble model named AIPStack has also been built but the AUC was only 0.8 and accuracy of 0.76 (Deng et al., 2022). Thus, there is a clear gap that remains and needs to be filled.

Moreover, there is a necessity for developing a new, more accurate prediction model which could both advance our knowledge of the relationship between peptide sequence and anti-inflammatory activity and serve as a reference for the rational design of AIPs that take into account the key elements provided by the model explanation. In order to design and validate the model for this investigation, we developed a new, larger dataset while keeping these concerns in mind.

Here, we have come up with an innovative approach of building a motif-analysis-driven (MAD) ML model, powered by datasets selected through multiple instances of random-sampling. The MAD-filtered datasets were used for training the model, including random forest, logistic regression, decision tree, Naive-Bayes, SVM and KNN. Exceptional performance was observed in case of the random forest model, which showed an accuracy of 99.17 and ROC of 99.75. The random forest model also displayed a specificity of 99.99% and a sensitivity of 95.16%. We tested our model practically by applying it to correctly identify a set of candidate AIPs which have already been experimentally-validated. 18 out of 19 experimentally-validated AIPs were identified correctly. We believe that our MAD-ML algorithm will be potentially useful for the AIP research. Finally, a web application was developed to host the MAD-ML-based AIP identification (AIPID) model. Further, all datasets used in building of this model were subjected to statistical analysis to extract features of AIPs and visually highight the core characteristics. All AIPs sourced from UniProt were deposited in the form of a repository onto our web application. The web application was finally named as AIPID.

## 2. Materials and methods

The pipeline of the AIPID model building consisted of several steps: 1. Collection of Positive and Negative dataset, 2. Motif analysis 3. Motif-analysis driven (MAD) dataset selection 4. Creation of a robust Classification model, 4. Creation of a Candidate dataset and its testing. Once the model was built, it was tested using the candidate dataset which contains experimentally-validated sequences.

Next, a web-based application was built to allow the users to easily access and utilize the MAD-ML-based AIPID facility for research purposes.

### 2.1 Data Fetching and Dataset creation

#### 2.1.1 Data fetching

The primary database for sourcing the AIPs includes UniProt, which was used specifically for fetching data for the creation of dataset (**UniProt Consortium et al., 2023**). Basic search filter was used for the collection of datapoints for both positive as well as negative datasets. For the collection of Anti-inflammatory peptides (Positive datapoints), keywords used to obtain the sequences were ‘Anti-inflammatory’, ’Antiinflammtory’. As a result, 1808 AIPs were found, all of which were used as the part of our positive dataset. For the collection of non-Anti-inflammatory peptides (Negative datapoints) keywords used were “NOT-Antiinflammatory”, “NOT-Anti-inflammatory” and the sequence length was kept in the range of 1-120 Ami no acids. A total of 39,009,166 (that is, more than 39 million peptides) were found to be anti-inflammatory.

#### 2.1.2 Dataset creation

All 1808 AIPs obtained from UniProt were included in our positive dataset (**UniProt Consortium et al., 2023**). We intended to take the ratio of positive:negative dataset as 1:5 and 1:10 for model building. Thus, it was not possible to include all 39,009,166 non-AIPs in our negative dataset. But to ensure that the unique features of all non-AIPs are represented in the negative dataset, we resolved to a custom algorithm that we call as MAD selection (motif-analysis driven selection). The main aim here was to choose the representative set of non-AIPs (9000 for maintaining the positive to negative dataset ratio of 1:5, and 18,000 for the 1:10 ratio). To select these representative datasets for non-AIPs (negative dataset), we first performed random sampling of the datasets to obtain 5 datasets of 18,000 AIPs each. We then performed motif analysis to compare all datasets and ensure that sequences that highlight all the top motifs are included in the final negative dataset. Redundancy removal was carefully ensured. The MAD-selection technique adopted to select the representative negative dataset (the most representative subset of the larger negative dataset), can be understood in detail from **Figure 1** below.

**Figure 1.**
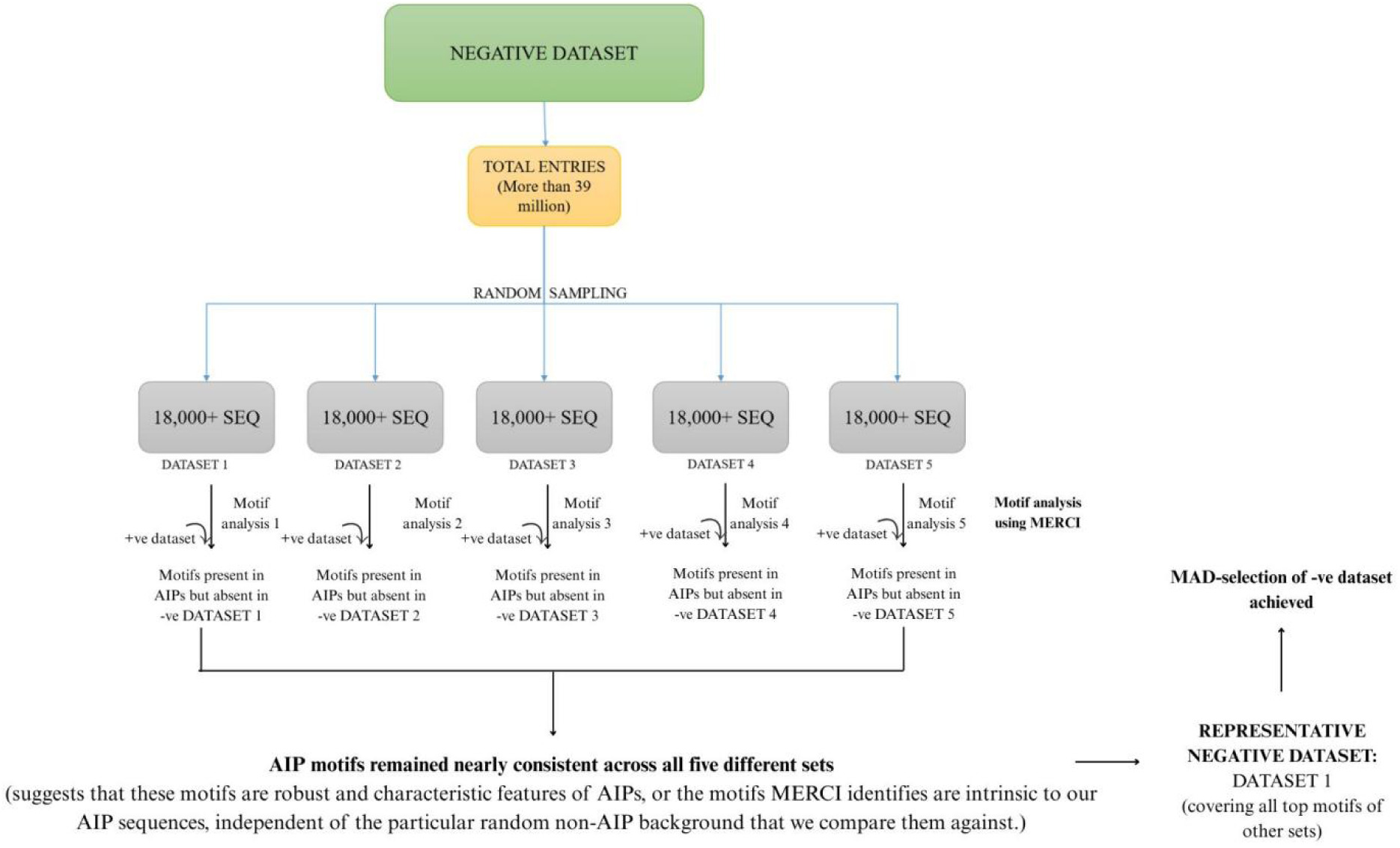
MAD-selection technique (or Motif-Analysis-Driven selection technique) for identification of the representative dataset for machine-learning model building

#### 2.1.3 Motif extraction and analysis using MERCI

For all 5 datasets, we extracted the motifs that are absent in a particular set of non-AIPs but present in AIPs, using the MERCI software. All 5 sets of non-AIPs were found to yield almost similar results. The motif analysis was done both in the absence as well as in the presence of outliers.

### 2.2 Feature Extraction

We used the ProtParam module from Biopython package (version 1.79) to calculate the peptide composition and physicochemical properties **(Gasteiger et al., 2005; Cock, et al.,2009)**. Propy3 Python package was utilized for calculating the CTD descriptors (https://github.com/MartinThoma/propy3). The ’composition’ features indicate the proportion of each type of amino acids in the peptide. The CTD descriptors describe peptide qualities such as ’hydrophobicity’, ’normalised van der Waals volume’, ’polarity’, and ’polarizability’, ‘charge’, ’secondary structure’ and ’solvent accessibility’.

### 2.3 Deploying Machine Learning Models

We built our predictive model using many of the algorithms which are primarily used for the binary classification. Our aim was to decide the best algorithm based on metrics results. All the models were used from ‘Scikit-learn’ python libraries **(Pedregosa et al., 2011)**. The data was split such that 20% represented the test data and 80% the train data. We tested models with different dataset ratios and different folds of cross-validation to achieve the best result.

#### 2.3.1 Random Forest

The Random Forest (RF) model is an ensemble prediction model that can do both regression and classification. RF has been used to categorise peptides and address several biological challenges **(Manavalan, et al., 2018)**. We compared the performance of RF with other classification techniques.

#### 2.3.2 Support Vector Machine

Support Vector Machine (SVM) is a popular classifier for peptide prediction **(Ng, et al., 2015)**. SVM is very effective for binary classification tasks. The model separates samples into classes using a hyperplane, which may be described in high-dimensional space using kernel transformations.

#### 2.3.3 K - nearest neighbour

The k-nearest neighbour (KNN) method is a supervised machine learning technique that is mostly used for classification applications. It has been extensively utilised for illness prediction. The KNN is a supervised algorithm that predicts the categorization of unlabelled data using the features and labels from the training data. Generally, the KNN method may classify datasets using a training model comparable to the testing question by considering the k nearest training data points (neighbours) that are closest to the query being tested. Finally, the algorithm uses the majority voting procedure to choose which categorization to finalise **(Uddin, et al., 2022)**.

#### 2.3.4 Naïve Bayes

Naive Bayes is a probabilistic classification technique that relies on Bayes’ theorem. It asserts independence among characteristics based on the class label, simplifying calculations. This method works particularly well for text categorization and spam filtering. Naive Bayes computes the probability of each class using the feature values and assigns the class with the highest probability to the input data point. Despite its simplicity and assumption of feature independence, Naive Bayes frequently performs quite well in reality and is computationally economical, making it a popular choice for numerous classification applications **(Haruechaiyasak, 2008)**.

#### 2.3.5 Decision tree

Decision trees are flexible and interpretable models for classification and regression. They recursively divided the dataset based on the most informative attributes, resulting in a tree-like structure. At each node, the decision tree chooses the feature that best separates the data based on parameters such as Gini impurity or information gain. The resultant tree is easily visualised, which aids interpretation. Decision Trees are robust against outliers, can handle both numerical and categorical data, and its ensembles such as Random Forests show improved predicted accuracy by mixing many trees.

#### 2.3.6 Logistic Regression

Logistic Regression is a linear classification approach that may be applied to both binary and multiclass issues. The logistic function is used to describe the chance that a data point belongs to a specific class. Training entails optimising coefficients to increase the probability of the observed data. Logistic Regression is frequently used in a variety of sectors because of its simplicity and interpretability.

### 4.4. Cross Validation

We conducted cross-validation using both five-fold and ten-fold methods on our training dataset. In the case of five-fold cross-validation, each fold consisted of 20% of the data, ensuring that the test sets contained 20% positive data and 20% negative data. Similarly, for ten-fold cross-validation, each fold comprised 10% of the data, resulting in test sets with 10% positive data and 10% negative data. To maintain class balance in the test sets, we employed stratified sampling.

All the ML algorithms were applied to three different models based on the ratio of positive to negative dataset and the folds of cross-validation, that is, Model 1 (1:5, 5 folds), Model 2 (1:5, 10 folds), and Model 3 (1:10, 5-folds).

### 4.5. Performance evaluation

We evaluated our models’ performance using common measures such as sensitivity, specificity, accuracy, Matthews’ correlation coefficient, ROC value and the harmonic mean of the precision-recall (F1) score.

**Sensitivity** is the metric that evaluates a model’s ability to predict true positives of each available category.

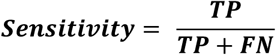

**Specificity** is the metric that evaluates a model’s ability to predict true negatives of each available category.

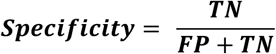

**Accuracy**, in machine learning, refers to the overall proportion of predictions the model makes that are correct. It’s a simple and intuitive metric to understand, but it has limitations, particularly in imbalanced datasets.

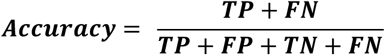

**F1 score** is the harmonic mean of precision and recall. The harmonic mean gives more weight to lower values, making it a stricter measure than the arithmetic mean (average). F-score is a metric used to evaluate the performance of a classification model, particularly in binary classification problems. It addresses the limitations of using just precision or recall alone by incorporating both metrics into a single score.

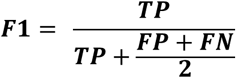

**MCC** is a more robust metric that considers both true and false positives/negatives. It takes into account the true negatives, which accuracy often overlooks. The range of MCC is between -1 and +1. A perfect score of +1 indicates perfect prediction, 0 represents a random guess, and values below 0 indicate worse-than-random performance.

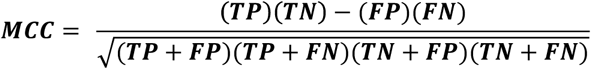

**ROC (Receiver Operating Characteristic)** is a curve that shows the trade-off between true positive rate (TPR) and false positive rate (FPR) for a classification model at different thresholds. The area under the ROC curve (AUC) is a measure of the overall performance of the model **(Hajian-Tilaki, 2013).**

### 4.6. Preparation of the candidate dataset

19 unique experimentally-validated AIPs freshly deposited on UniProt (which were initially unavailable) were downloaded at the time of testing the final model and were subjected to the algorithm to determine the efficacy of the model.

### 4.7 App Development and Deployment

To ensure user-friendly accessibility of the MAD-ML-based AIPID prediction model to researchers, clinicians, and bioinformaticians without programming expertise, a dedicated web-based application was developed. The platform was designed to allow users to input peptide sequences, retrieve anti-inflammatory activity predictions, and explore curated peptide datasets interactively through an intuitive graphical user interface.

#### 4.7.1 Backend Development

The Random Forest machine learning model, which demonstrated superior performance among all evaluated algorithms (Section 6.2), was serialized using Python’s pickle module and stored as a .pkl file for deployment. Real-time peptide feature extraction modules were implemented using Biopython (v1.83) and Propy3, enabling the calculation of sequence-derived physicochemical properties and composition-based descriptors. All peptides used for model training and evaluation were sourced exclusively from the UniProt database. Further, these datasets were subjected to statistical analysis to identify and visualize core sequence characteristics of anti-inflammatory peptides (AIPs), which guided feature selection and model refinement.

#### 4.7.2 Frontend Development and Deployment

The frontend of the web application was developed using the Streamlit (v1.33.0) framework, leveraging its multi-page app feature to organize dedicated sections for prediction, dataset browsing, statistical visualizations, help documentation, contact details, and developer profiles. Key Python packages utilized in the frontend development included streamlit, scikit-learn, biopython, propy3, pandas, numpy, altair, openpyxl, and setuptools for managing data display, interactive plotting, and backend-frontend integration.

The application was deployed via Streamlit Community Cloud (https://streamlit.io/cloud) by linking a publicly accessible GitHub repository containing the trained model, pages and a custom requirements.txt file specifying all package dependencies. The final web application, titled AIPID, integrates the motif-analysis-driven AIPID prediction engine and a searchable repository of UniProt-derived AIP sequences, enabling seamless identification, analysis, and visualization of anti-inflammatory peptide candidates. The publicly available application can be accessed at: https://aipid-app-version1.streamlit.app/.

## 3. Results and Discussion

Our model was able to successfully predict peptides active against inflammation.

### 3.1 Characteristics of positive dataset

#### 5.1.1. Sequence length

Our positive dataset initially consisted of 1808 data points but after pre-processing we removed the sequences with unknown character ‘X’ which led to a total of 1785 sequences. The sequence length of our positive data points ranged from 10 to 4578, but the median length of our dataset was found to be 260 amino acids. The distribution of sequence length in the positive dataset, both in the absence as well as the presence of outliers can be seen in **Figure 2**.

**Figure 2.**
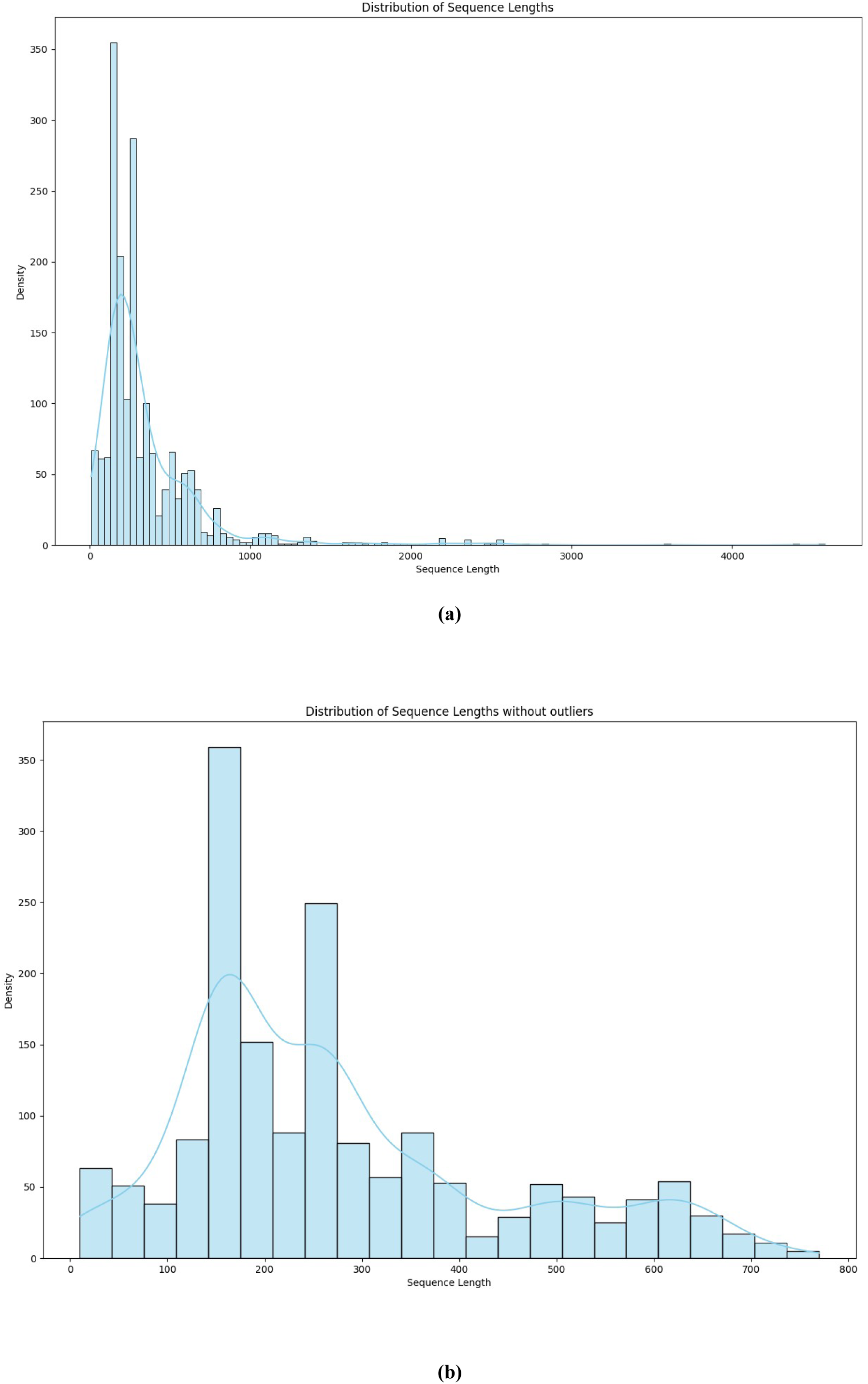
Distribution of sequence of length of AIPs. (a) In presence of outliers (b) In absence of utliers.

#### 5.1.2. Amino acid composition

In our positive dataset it was seen that leucine has the highest percentage in total dataset followed by serine and alanine. Tryptophan was found to contribute the least to our sequences. The contribution of amino acids in our positive dataset has been shown in **Figure 3** **(a).**

**Figure 3.**
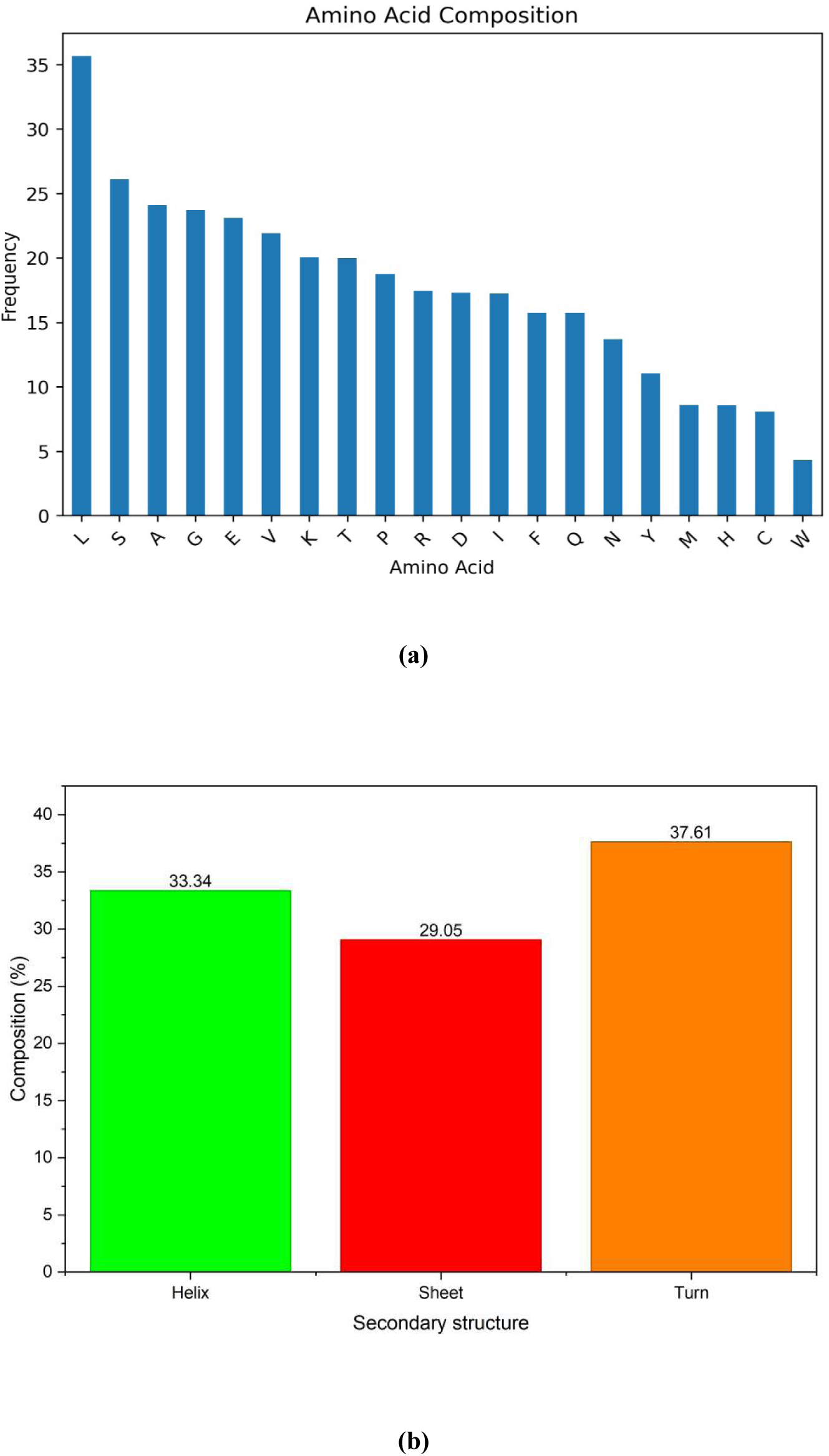
Amino acids and secondary structure composition of AIPs. (a) Contribution of different amino acids to the AIPs (b) Composition of secondary structure fractions of AIPs

#### 5.1.3. Secondary Structure Analysis

We used the secondary structure composition as a feature. It was seen that sheet was in majority with 37.61% in the total dataset followed by helix and turn which were 33.34% and 29.04% respectively. The composition of secondary structure fractions has been shown in **Figure 3** **(b).**

### 3.2 Results from motif analysis of AIPs validates our random sampling strategy

The motif analysis for AIPs was performed separately for the 5 different dataset combinations (due to the fact that the positive was same but negative dataset was of five types). The AIP motifs remained consistent across all five different sets, which suggests that these motifs are robust and characteristic features of AIPs. In other words, the motifs MERCI identifies are intrinsic to our AIP sequences, independent of the particular random non-AIP background that we compare them against. Additionally, this also means that the ‘random negatives’ were appropriate. It also indicates that our negative datasets (non-AIPs) were random and sufficiently diverse and that none of them inadvertently contained significant numbers of AIP-like sequences that might otherwise obscure or distort motif identification. This validates our random sampling strategy for negatives.

It can be said that the motif signal of AIPs is stronger than noise, meaning that the motifs found are likely strong discriminators of AIPs because even when the “noise” of the negative background changes slightly (across 5 different random non-AIP sets), the motifs remain largely the same.

Presence/absence of outliers had negligible impact on motif results. Since motif patterns persisted both in the presence and absence of outliers, it suggests that our AIP motif signature is strongly conserved and not driven by a few extreme sequences.

PGWF, PGWFL, GWFLC, PGWFLC, FVNVT, FVNVTD and FSESAA are some of the top motifs occurring in AIPs which are absent in non-AIPs. Among the five negative datasets, Negative Dataset 1 was selected as the representative dataset, as it yielded the majority of the characteristic motifs identified in our analysis. The top motifs present in AIPs but absent in non-AIPs present in Negative Dataset 1 can be studied in detail in **Figure 4**. Similar results were obtained when other negative datasets were used. These can be found in **supplementary file** as **Figure S1-S4.**

**Figure 4.**
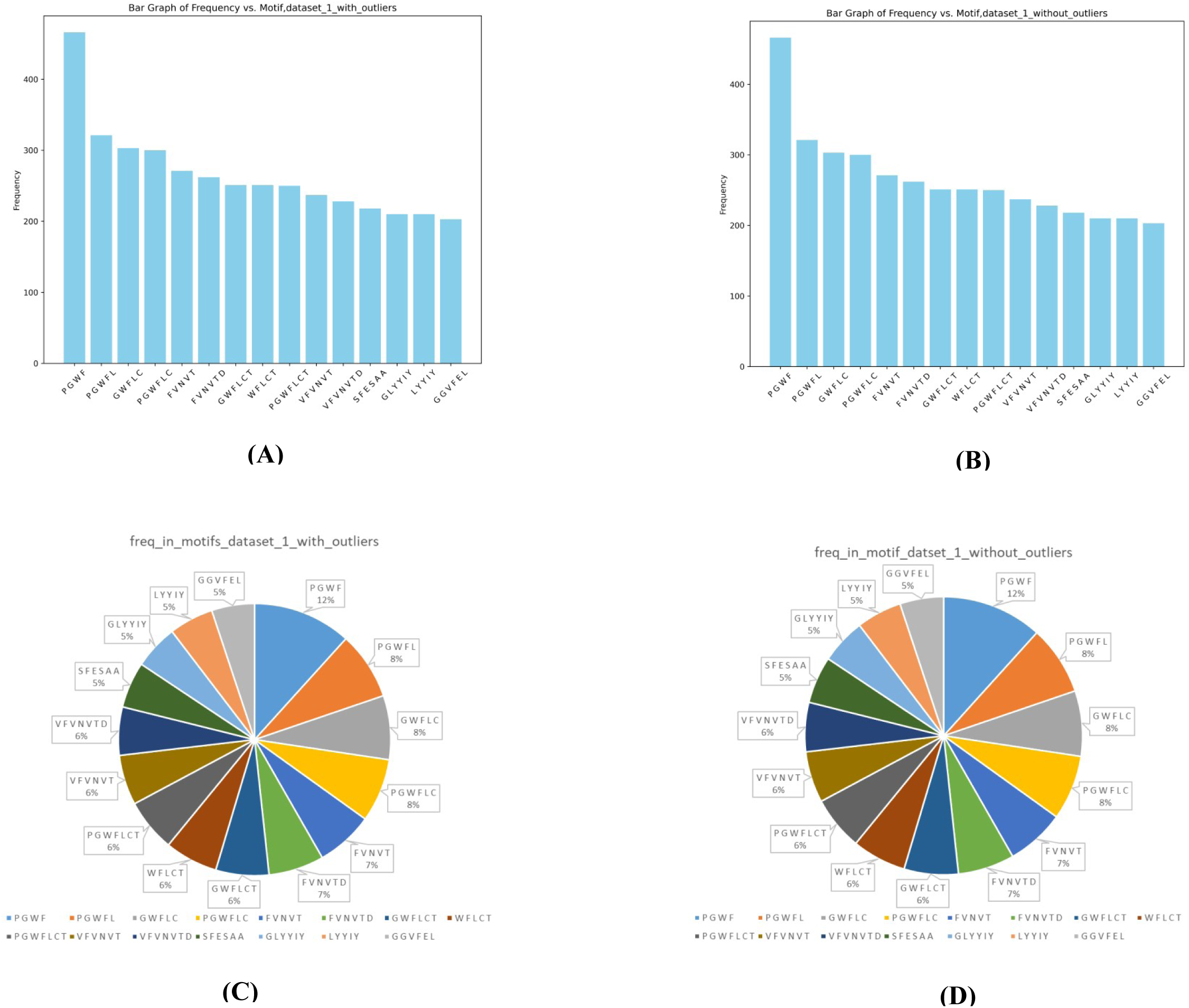
Motifs absent in non-AIPs (representative negative-dataset-1) but present in AIPs

### 3.3 Model creation, training and evaluation

The original positive dataset and the representative negative-dataset-1 (which represented all other negative datasets) were selected for training the model.

After the anti-inflammatory peptide dataset was well characterised, we utilised the peptides’ primary and secondary structural data to create machine learning models that could recognise and comprehend characteristics that could be particular to AIPs. We trained our machine learning models and employed a total of 175 features that came from CTD analysis and basics parameters.

The objective was to identify models that not only minimize the rates of false positive but also perform well across other metrics.

The random forest performed well across all the combinations, achieving great metrics. The scores for all the models at different folds of cross-validation have been summarized in **Table 1**. The same can be visualized in **Figure 5** for a more comprehensive understanding. The random forest model produced a very low number of false positives. There was found to be only 1 false positive in the 1:5 dataset ratio for both 5 folds and 10 folds cross validation configuration, and only 2 false positives for the 1:10, 5 fold scenario. The model wasn’t just good at avoiding the false negative, but also performed consistent across all other metrics with f1 score above 0.95, indicating its excellent precision and recall balance also roc over 0.94, demonstrating true positives relative to the false positive rate.

**Table 1.** Performance evaluation scores for all models

| Model 1 (1:5,5) | FP | FN | TP | TN | SN | SP | Accuracy | AUC | F1 |
| --- | --- | --- | --- | --- | --- | --- | --- | --- | --- |
| Logistic Regression | 16 | 220 | 1208 | 7123 | 84.59 | 99.77 | 97.2 | 0.98 | 0.93 |
| Naïve Bayes | 8 | 120 | 1308 | 7131 | 91.59 | 99.88 | 98.5 | 0.96 | 0.95 |
| **Decision Tree** | 71 | 37 | 1391 | 7068 | 97.4 | 99 | 99.7 | 0.98 | 0.97 |
| **Random Forest** | 1 | 69 | 1359 | 7138 | 95.16 | 99.98 | 99.2 | 0.99 | 0.98 |
| **SVM** | 0 | 208 | 1220 | 7139 | 85.54 | 1 | 97.6 | 0.97 | 0.93 |
| **KNN** | 852 | 105 | 1323 | 6287 | 92.64 | 88.06 | 88.8 | 0.96 | 0.76 |
| **Model 2 (1:5,10)** | **FP** | **FN** | **TP** | **TN** | **SN** | **SP** | **Accuracy** | **AUC** | **F1** |
| **Logistic Regression** | 19 | 211 | 1217 | 7120 | 85.22 | 99.73 | 97.3 | 0.98 | 0.93 |
| **Naïve Bayes** | 6 | 119 | 1309 | 7133 | 91.66 | 99.91 | 98.5 | 0.96 | 0.95 |
| **Decision Tree** | 79 | 40 | 1388 | 7060 | 97.19 | 98.89 | 98.7 | 0.98 | 0.96 |
| **Random Forest** | 1 | 70 | 1358 | 7138 | 95.09 | 99.98 | 99.2 | 0.99 | 0.98 |
| **SVM** | 1 | 204 | 1224 | 7138 | 85.71 | 99.99 | 97.6 | 0.97 | 0.93 |
| **KNN** | 812 | 99 | 1329 | 6327 | 93.06 | 88.62 | 89.4 | 0.96 | 0.76 |
| **Model 3 (1:10,5)** | **FP** | **FN** | **TP** | **TN** | **SN** | **SP** | **Accuracy** | **AUC** | **F1** |
| **Logistic Regression** | 11 | 284 | 1144 | 14362 | 0.8 | 0.99 | 98.1 | 0.96 | 0.87 |
| **Naïve Bayes** | 85 | 141 | 1287 | 14288 | 0.9 | 0.99 | 98.6 | 0.94 | 0.91 |
| **Decision Tree** | 110 | 76 | 1352 | 14263 | 0.94 | 0.99 | 99 | 0.98 | 0.92 |
| **Random Forest** | 2 | 84 | 1344 | 14371 | 0.94 | 0.99 | 99.4 | 0.94 | 0.95 |
| **SVM** | 2 | 272 | 1156 | 14371 | 0.8 | 0.99 | 98.3 | 0.94 | 0.88 |
| **KNN** | 898 | 117 | 1311 | 13475 | 0.91 | 0.93 | 93.6 | 0.96 | 0.72 |

**Figure 5.**
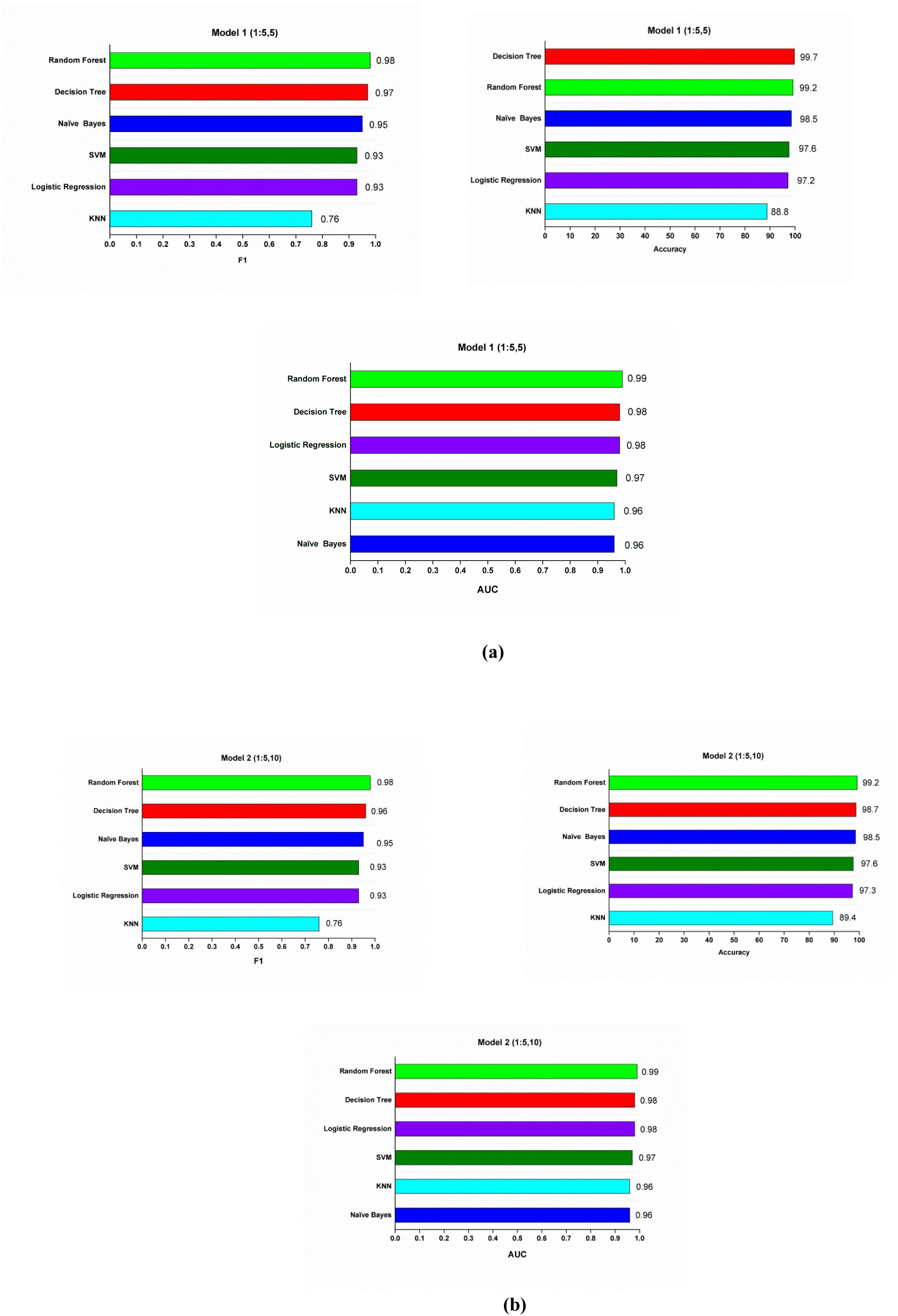
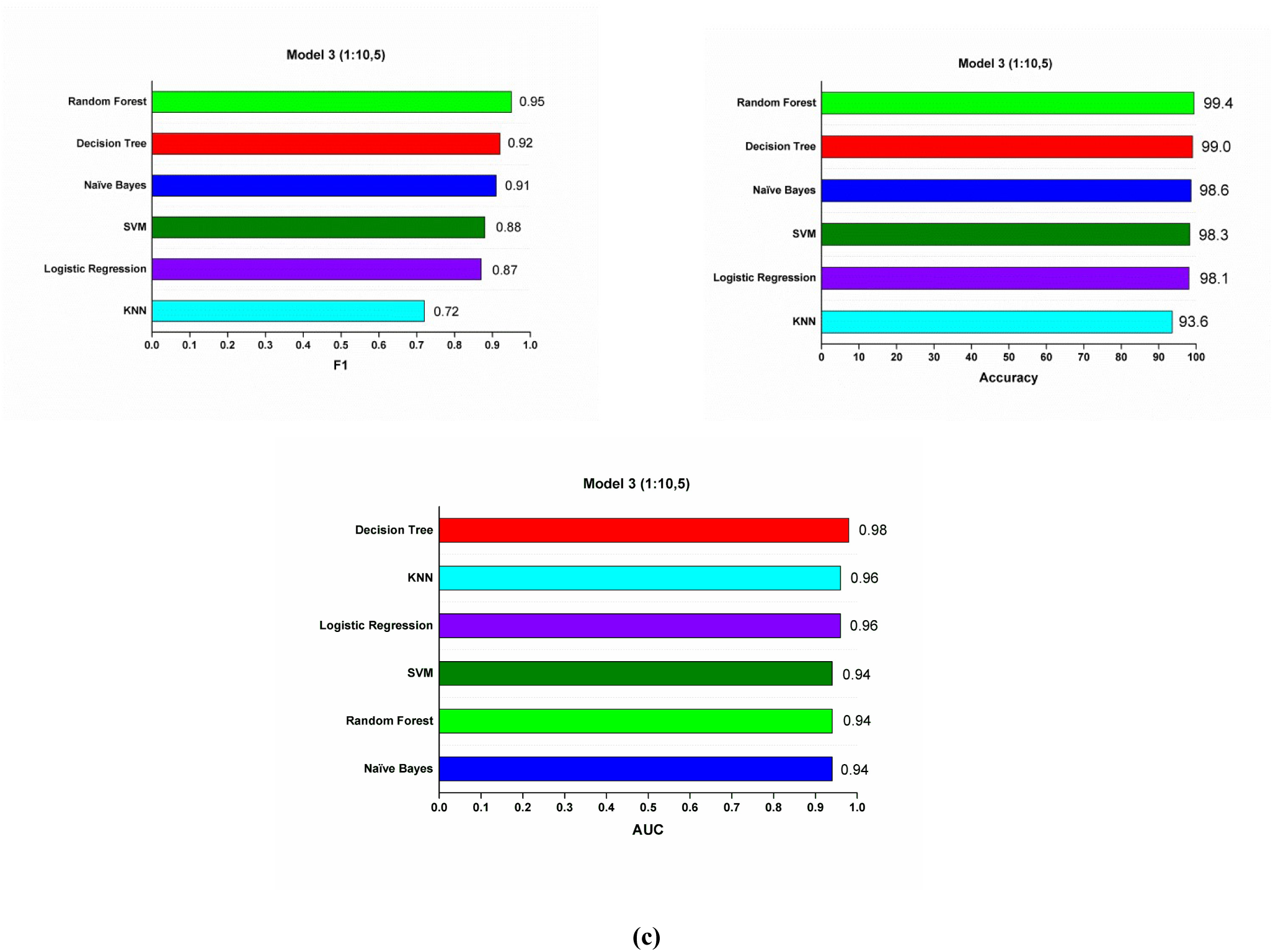
Metrics evaluation for 3 different models varying on basis of cross-fold of validation and ratio of positive and negative datasets. (a) Model 1 with positive:negative dataset ratio equal to 1:5 and 5-fold cross-validation (b) Model 2 with positive:negative dataset ratio equal to 1:5 and 10-fold cross-validation (c) Model 3 with positive:negative dataset ratio equal to 1:10 and 5-fold cross-validation

The Support Vector Machine (SVM) performed well in the dataset 1:5 dataset, where the false positive was as low as Random forest. It maintained a high F1 score of 0.93 and ROC above 0.97.

The decision tree classifier, which can be considered as the lower counterpart of Random Forest, performed considerably well with false positive being lowest at 110 in the 1:10, 5-fold dataset. It held a strong F1 score of 0.92, suggesting a good model.

The Naïve Bayes also performed well in some context, with false positive as low as 6 in 1:5,10 folds dataset. It showed increased false positive in the 1:10 dataset, while maintained a F1 score near 0.91.

Logistic Regression being a balance in complexity and performance, performs very well but not gaining the lows of false positive set by random forest or SVM, with its false positive not exceeding 19 and F1 score of 0.93.

Contrastingly, the KNN model had the highest occurrence of false positive i.e. 898 in the 1:10,5-folds, Other metrics also underperformed compared to the other models.

In conclusion, the random forest algorithm with its ensemble approach proved to be the best algorithm for our model offering a low false positive in the configuration of one 1:5 dataset, 10 folds. The model achieved a very low false positive of 1 and false negative very low of 69 in 1: 5 ratio 5 folds and 70 at 1:5 ratio 10 folds. The robustness of this model can be also validated by its high F1 score exceeding 0.95 and ROC about 0.94. The metrics achieved signify its reliability and predictive accuracy given its achievement in both minimisation and other performance parameters. The Random Forest model with 1:5 ratio of dataset and 5-fold cross-validation was used as our final predictive model.

### 3.4 Model testing using candidate dataset

Remarkably, 18 out of 19 experimentally-validated AIPs in our candidate dataset were correctly identified as positives using the random forest model (1:5, 5-fold). The details of the candidate dataset and the predictions made by our MAD-ML AIPID model have been dsiscussed in **Table 2**.

**Table 2.**
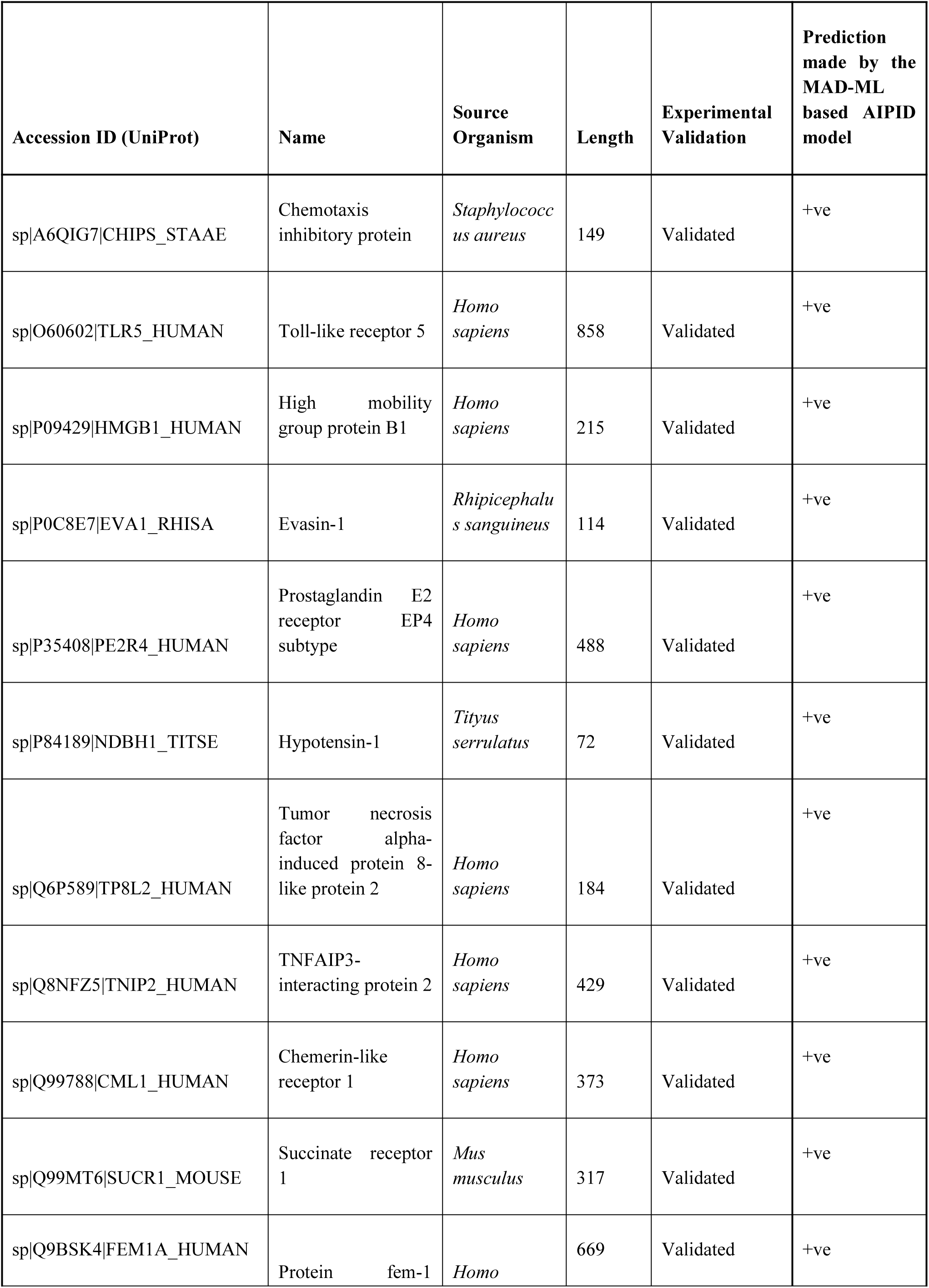
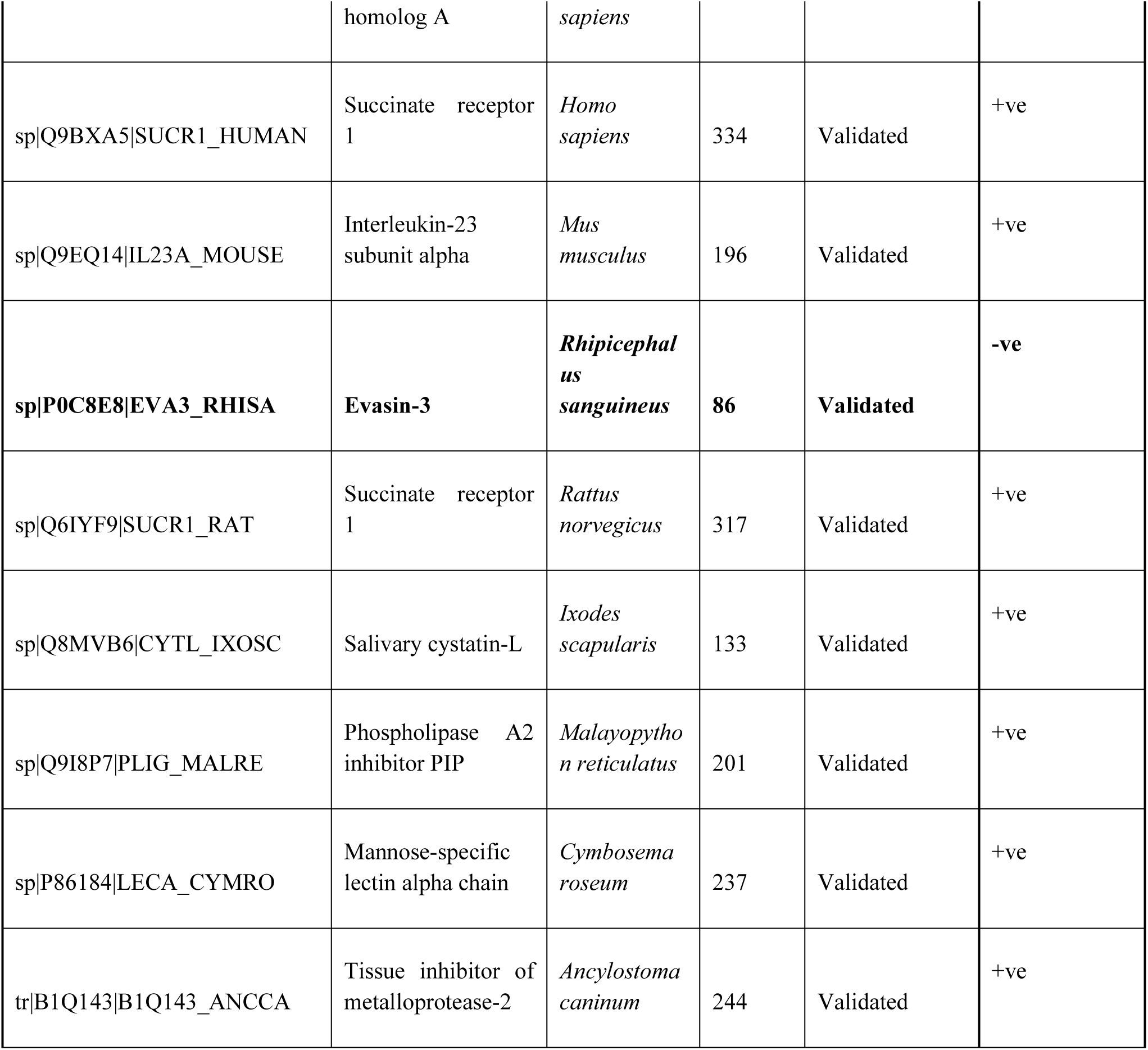
Details of the candidate dataset used for performance evaluation and the predictions made by MAD-ML AIPID

### 3.5 Comparison with existing models

Models demonstrate impressive outcomes but still fall short. Among the stated approaches, our model stands out for its precision and accuracy, as seen by its exceptional performance numbers. With sensitivity (SN) of 95.23% and specificity (SP) of 99.98%, it outperforms other models in recognizing both true positives and true negatives. Its Matthew’s Correlation Coefficient (MCC) of 0.98 and Accuracy (ACC) of 99.34% demonstrate its strong prediction powers and remarkable balance of sensitivity and specificity.

An Area Under the ROC Curve (AUC) value of 1 indicates an optimal model with a flawless classification rate. This is a substantial improvement over previous models such as AIPpred and PreAIP 2019, which perform poorly overall. Notably, the model by Khan et al., 2023 and the AIPs-SnTCN model demonstrate remarkable results but still fall short. The performance scores for the existing models in comparison with our model have been stated in **Table 3**.

**Table 3.** Performance evaluation metrics for our model along with other existing models

| Method | SN (%) | SP (%) | MCC | ACC (%) | AUC | References |
| --- | --- | --- | --- | --- | --- | --- |
| AIPpred | 75.8 | 71.1 | 0.46 | 73 | 0.801 | Manavalan et al., 2018 |
| PreAIP | 63.2 | 90.3 | 0.566 | 79.5 | 0.833 | Khatun et al., 2019 |
| Optimized Feature Learning for Anti-Inflammatory Peptide Prediction Using Parallel Distributed Computing | 89.01 | 82.35 | 0.711 | 83.4 | NA | Khan et al., 2023 |
| AIPs-SnTCN | NA | NA | NA | 95.86 | 0.97 | Raza et al., 2023 |
| AIP Stack 2022 | NA | NA | 0.51 | 75.5 | 0.819 | Deng et al., 2022 |
| Prediction of anti-inflammatory proteins/peptides: an insilico approach | NA | NA | 0.58 | 78.1 | NA | Gupta et al., 2017 |
| AIPID | 95.23 | 99.98 | 0.98 | 99.34 | 1 | Anurag Anand and Urkude et al., 2024 |

## 4. AIPID Web Application Interface and Performance

To enhance the accessibility of the MAD-ML-based AIPID prediction model for researchers, clinicians, and bioinformaticians, a dedicated standalone web platform, titled AIPID, was developed. The application is designed to accept peptide sequences in single-letter amino acid code format and deliver real-time predictions regarding their anti-inflammatory potential.

The platform’s interface is organized via a sidebar-driven multi-page layout developed using the Streamlit (v1.33.0) framework. The Home page serves as the central prediction module powered by the integrated AIPID engine. Users can input peptide sequences and receive classification outcomes alongside a model-derived confidence score generated from the Random Forest classifier’s probability estimates.

In addition to the prediction module, AIPID includes several dedicated pages, like AIP Features and others. AIP Features is a searchable and browsable repository containing anti-inflammatory peptides sourced from the UniProt database, accompanied by basic statistical summaries and sequence-based descriptors. The About, Help, Contact, and Developer Information sections provide platform background, user instructions, correspondence details, and contributor profiles respectively.

The application processes user-submitted sequences in real time through integrated Biopython and Propy3-based feature extraction pipelines, which compute essential sequence-derived physicochemical properties and composition-based descriptors. Predictions are performed using the trained Random Forest model, serialized as a .pkl file and deployed within the backend. The AIPID platform demonstrated consistent and reliable performance in classifying both validated anti-inflammatory peptides and synthetic sequences, achieving accurate predictions with minimal latency. The integrated statistical analysis functionality provides additional insight into characteristic features of anti-inflammatory peptides, aiding users in sequence interpretation and evaluation.

The complete application was publicly deployed via Streamlit Community Cloud (https://streamlit.io/cloud) and is accessible freely at https://aipid-app-version1.streamlit.app/ , with the trained model, datasets, and deployment configuration files maintained within a public GitHub repository (https://github.com/AnanyaAnuragAnand/AIPID-app/).

A screenshot of the Home page featuring the deployed AIPID prediction tool within AIPID is presented in **Figure 6**. **Figure 7** illustrates the statistical visualization section available on the AIP Features page.

**Figure 6.**
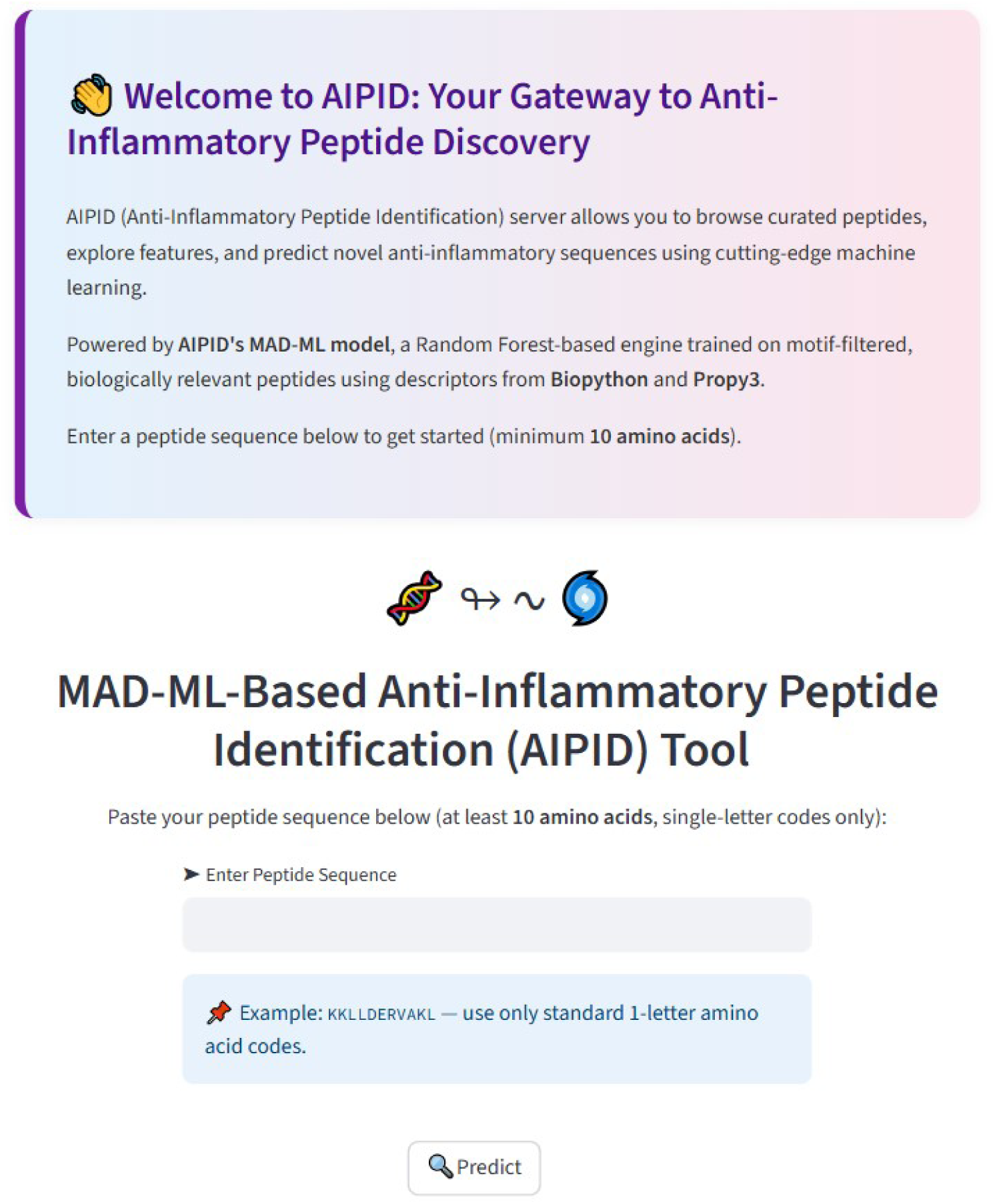
App page of AIPID hosting the MAD-ML AIPID model. The input field is designed for evaluating sequences with 10 or more amino acids. The predict button helps the user to submit the query sequence and, thereby, obtain the prediction results.

**Figure 7.**
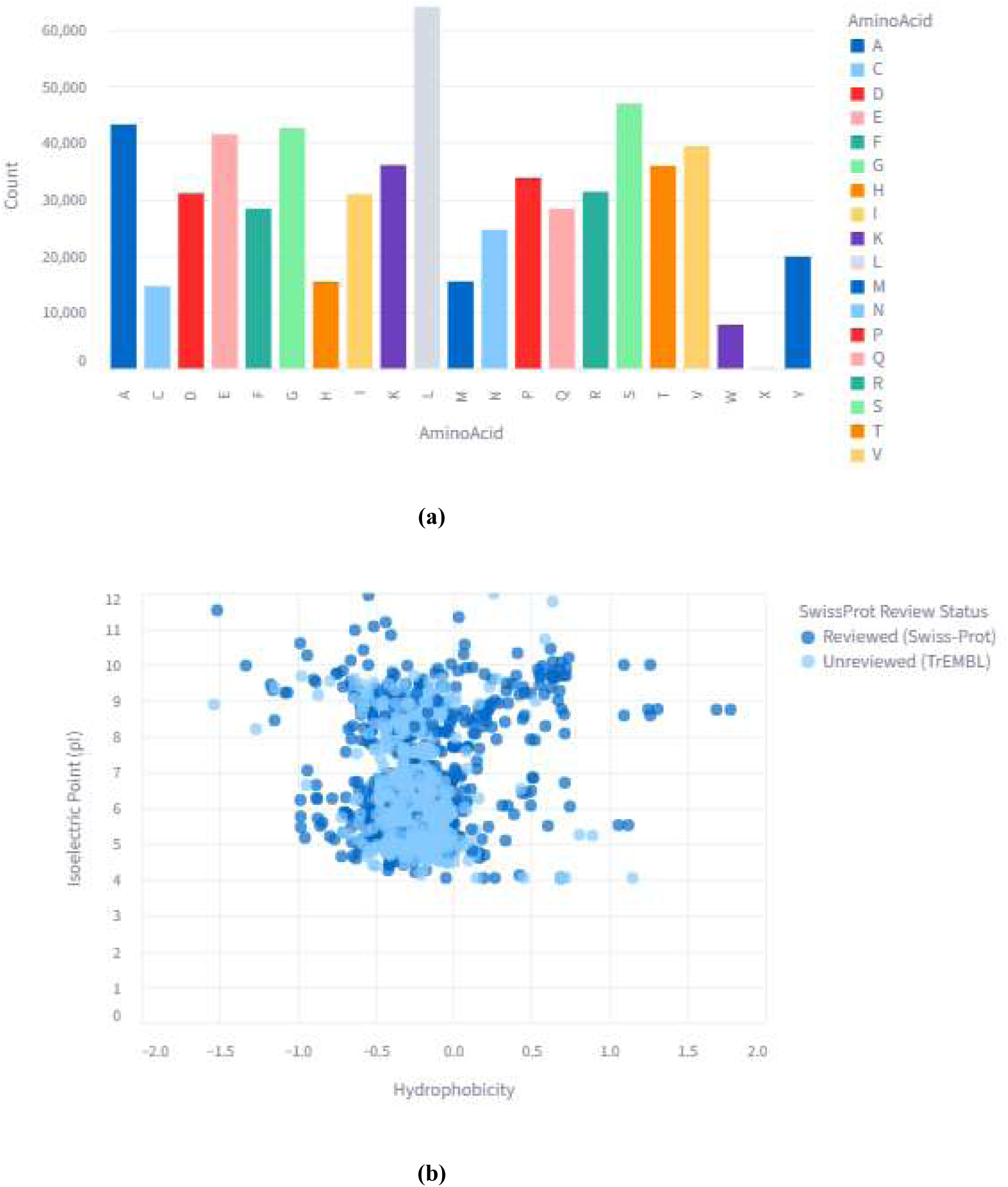
Statistics displayed on the AIP Features page of the AIPID application. (a) Amino acid distribution across all AIPs (b) Plot showing pI and hydrophobicity measurements for all AIPs

## 5. Conclusion and Future Perspectives

In this study, we successfully developed a motif-analysis-driven machine learning model for the prediction of anti-inflammatory peptides (AIPs). Among the models evaluated, the Random Forest classifier demonstrated the best performance, achieving a sensitivity of 95.16%, specificity of 99.98%, and an F1 score of 98.14%. The model’s ability to correctly predict 18 out of 19 experimentally validated AIPs highlights its robustness and practical applicability for peptide-based therapeutic discovery.

To ensure broader accessibility and usability, we integrated the trained model into a publicly available, interactive web platform named AIPID. This platform not only hosts the AIPID prediction tool but also includes a searchable repository of UniProt-derived AIP sequences, along with essential statistical insights. AIPID provides an intuitive, multi-page interface for researchers, clinicians, and students to explore, predict, and analyze anti-inflammatory peptide sequences without requiring prior computational expertise.

Looking ahead, future developments shall focus on expanding the AIP dataset to incorporate newly identified experimental sequences, integrating additional physicochemical and structural descriptors, and extending the platform’s capabilities for de novo peptide design and immunomodulatory activity prediction. Furthermore, incorporating deep learning architectures and ensemble modeling strategies could enhance prediction accuracy and broaden the platform’s scope to include peptides with other immunotherapeutic properties.

Finally, we would like to highlight that the motif-analysis-driven (MAD) selection strategy introduced in this study represents a significant methodological advancement for peptide-based machine learning applications. By integrating iterative random sampling with motif profiling, the MAD-ML technique ensures the selection of representative, diverse, and non-redundant negative datasets, addressing a critical challenge in imbalanced classification problems, particularly in bioinformatics where comprehensive negative datasets are scarce or overwhelming in size. The success of this strategy in achieving superior predictive performance for anti-inflammatory peptide identification highlights its potential as a broadly applicable framework for future peptide classification models. Beyond anti-inflammatory peptides, the MAD-ML approach can be adapted for the development of predictive models targeting other bioactive peptide classes such as antimicrobial, anticancer, antihypertensive, or antiviral peptides. Its utility lies in systematically capturing sequence diversity while preserving informative motifs, thus improving model robustness and generalizability. Future work may focus on extending MAD-ML to incorporate structural motifs, post-translational modifications, or evolutionary conservation patterns, further enhancing its predictive capabilities. Additionally, integrating the MAD-ML selection pipeline with advanced deep learning architectures could open new avenues for high-accuracy, motif-informed peptide function prediction. We anticipate that the MAD-ML framework will serve as a valuable methodological asset for the peptide bioinformatics community, aiding in the systematic development of balanced, motif-rich training datasets for a variety of therapeutic peptide discovery pipelines.

## 6. Data availability

The source code, trained machine learning model, and deployment files for the AIPID platform are publicly available via our GitHub repository at https://github.com/AnanyaAnuragAnand/AIPID-app. The repository includes the complete Streamlit application, feature extraction modules, and the serialized Random Forest model (aipid_model.pkl) used for anti-inflammatory peptide prediction.

The original datasets used for model training and evaluation, comprising positive (anti-inflammatory peptides), negative (non-anti-inflammatory peptides), and candidate sequences, were sourced exclusively from the UniProt database. Due to data licensing policies and to avoid redundancy, these datasets are not separately hosted within the GitHub repository. However, users can reconstruct the datasets by retrieving the respective UniProt accession numbers and sequences, as described in the Methods section of this manuscript.

Detailed instructions for model use, application deployment, and dataset reconstruction are provided within the README.md file of the GitHub repository. The publicly accessible web application, AIPID, can be accessed at https://aipid-app-version1.streamlit.app/ , providing interactive prediction and exploration of anti-inflammatory peptide sequences.

## Supporting information

Supplementary_file_Anand_and_Urkude_et_al

