## Supplementary_file_Anand_and_Urkude_et_al for "AIPID: MAD-ML-Powered AIP Discovery Platform"

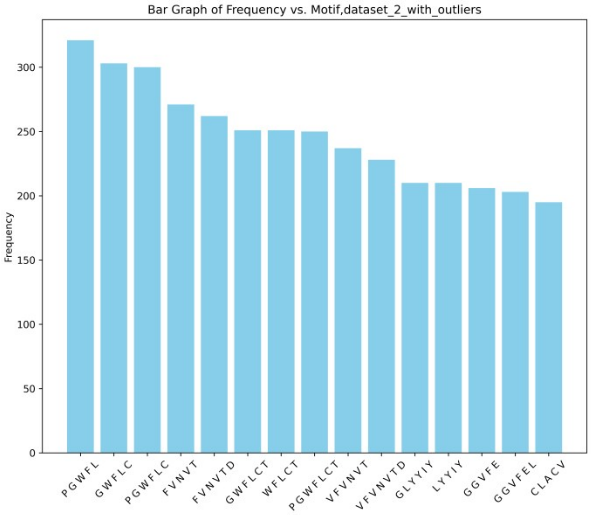


**(A)**

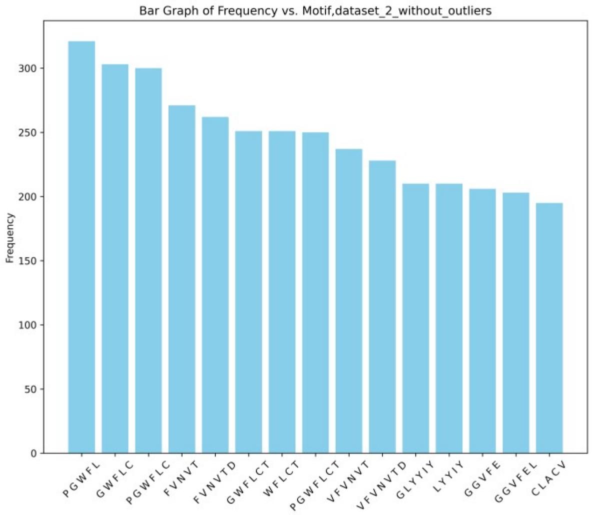


**(B)**

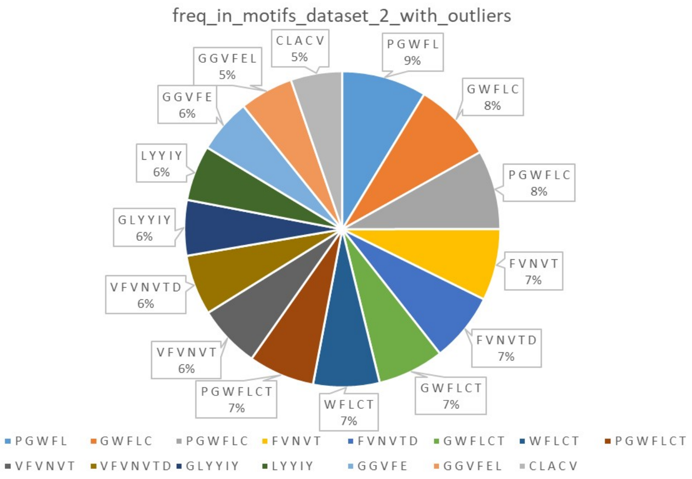


**(C)**

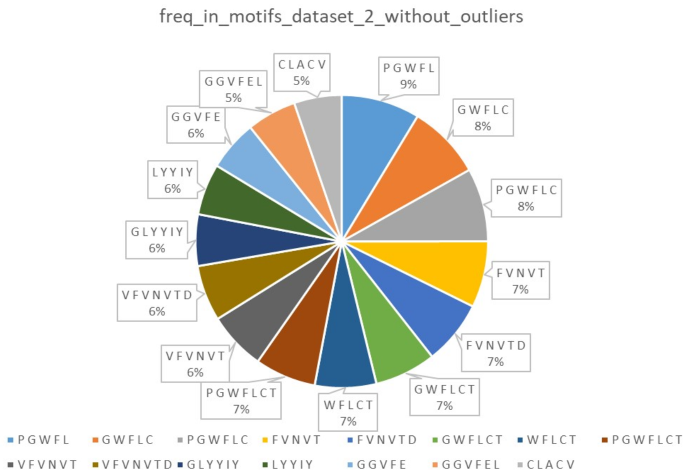


**(D)**

Figure S1. Motifs absent in non-AIPs (dataset-2) but present in AIPs

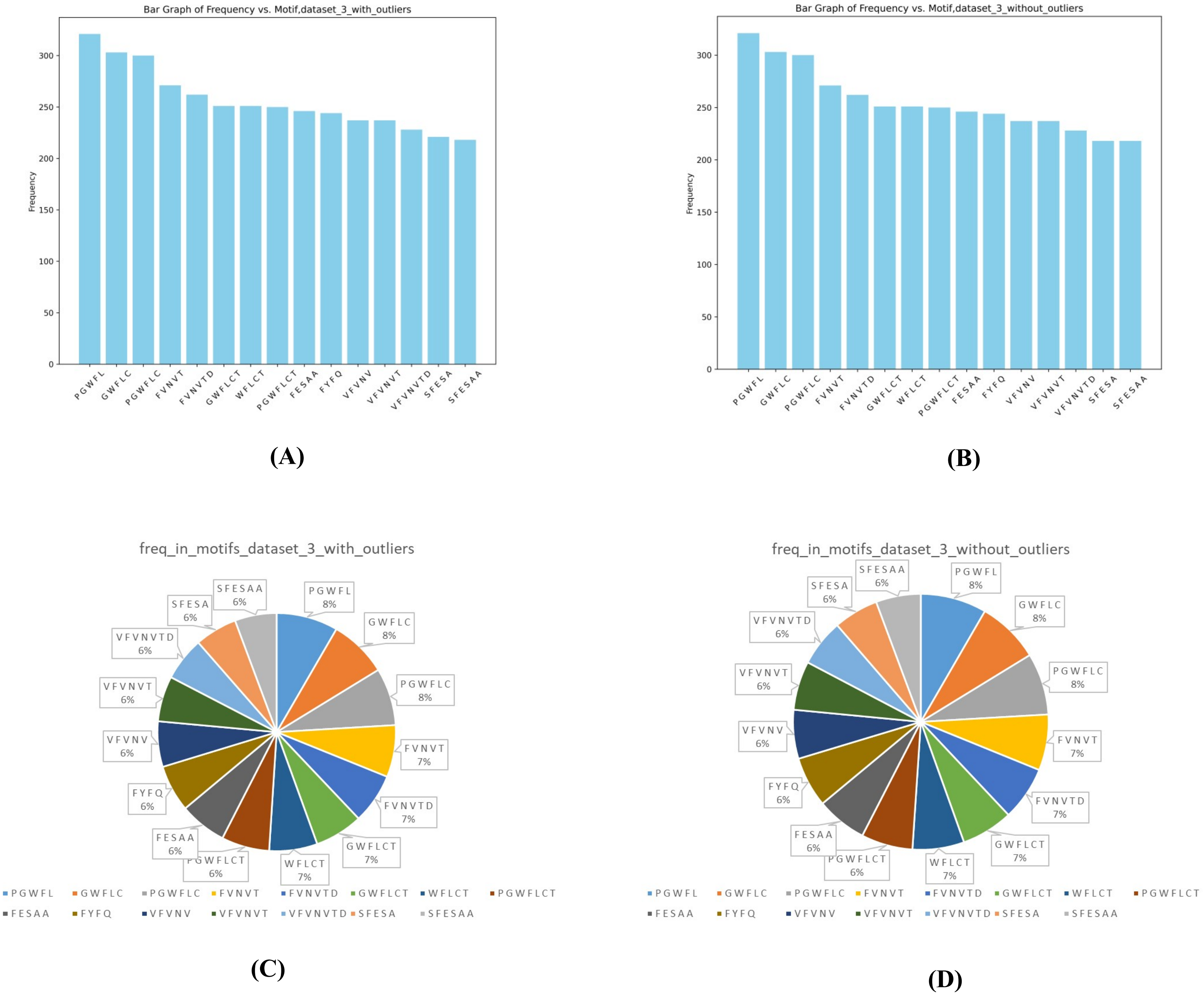


Figure S2. Motifs absent in non-AIPs (dataset-3) but present in AIPs

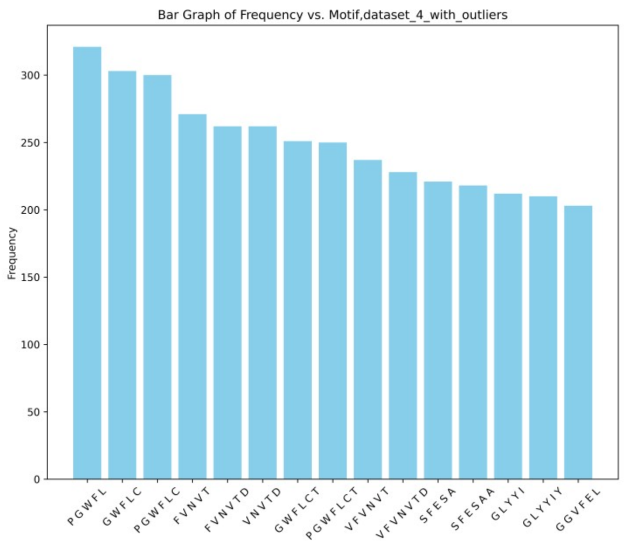


**(A)**

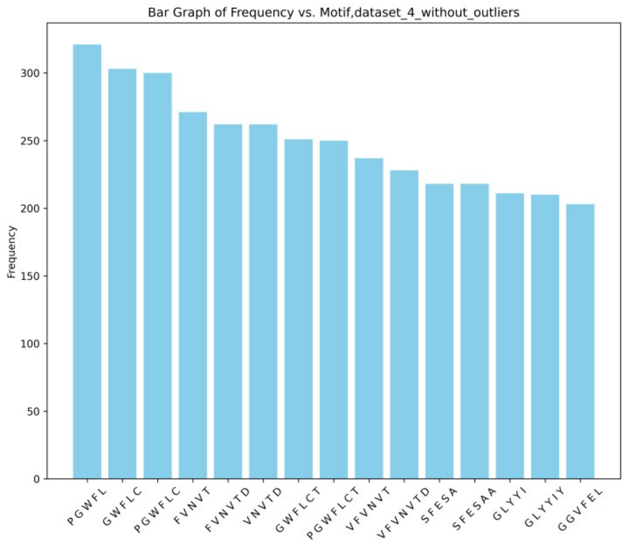


**(B)**

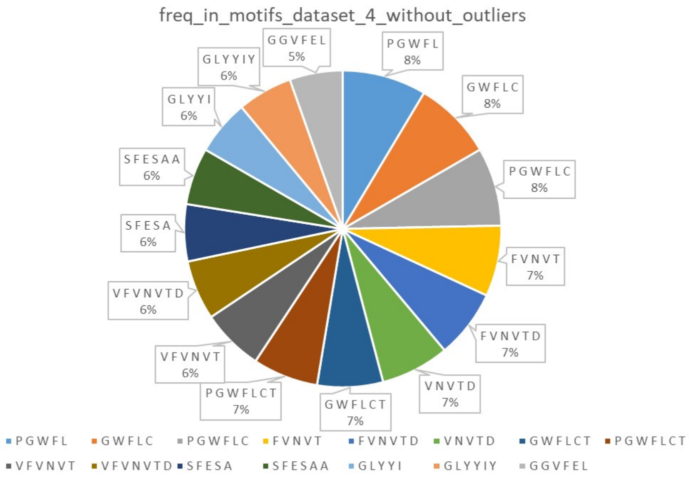


**(C)**

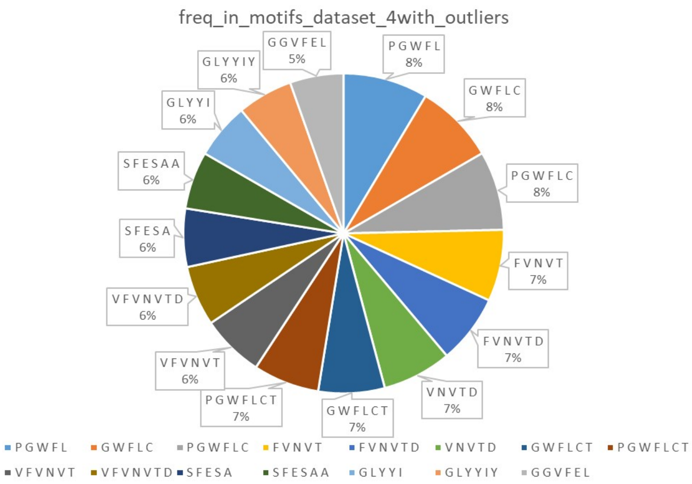


**(D)**

Figure S3. Motifs absent in non-AIPs (dataset-4) but present in AIPs.

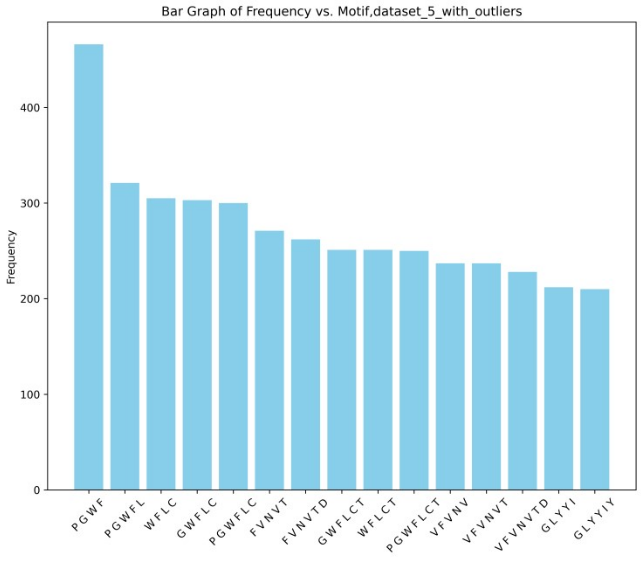


**(A)**

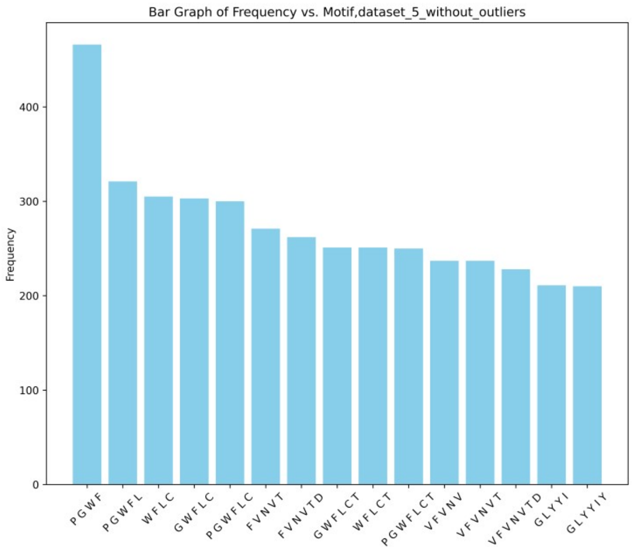


**(B)**

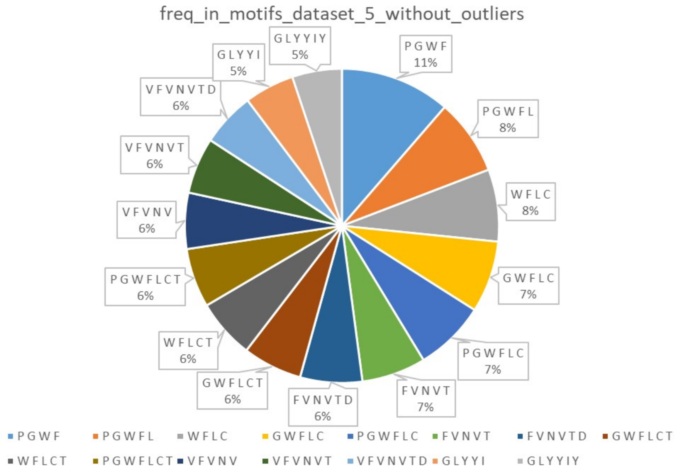


**(C)**

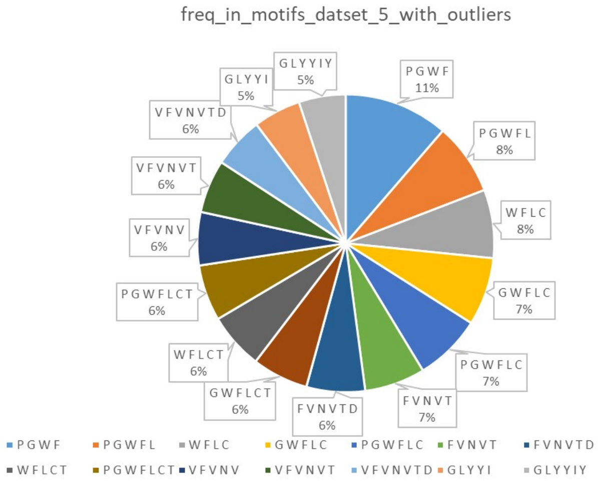


**(D)**

Figure S4. Motifs absent in non-AIPs (dataset-5) but present in AIPs.
